# Revisiting the origin of interleukin 1 (IL-1) based on biological activities of IL-1 in anamniotes and their sub-functionalization in amniotes

**DOI:** 10.1101/2022.02.15.480490

**Authors:** Eva Hasel de Carvalho, Eva Bartok, Helen Stölting, Baubak Bajoghli, Maria Leptin

**Affiliations:** European Molecular Biology Laboratory (EMBL), Directors’ Research. Meyerhofstrasse 1, 69117-Heidelberg, Germany; Institute of Clinical Chemistry and Clinical Pharmacology, University Hospital, University of Bonn. Venusberg Campus 1, 53127 Bonn, Germany; Unit of Experimental Immunology, Institute of Tropical Medicine, 2000 Antwerp, Belgium; National Heart and Lung Institute, Faculty of Medicine, Imperial College London, London, United Kingdom; Department of Hematology, Oncology, Immunology, and Rheumatology, University Hospital of Tübingen, Otfried-Müller-Strasse 10, 72076-Tübingen, Germany

## Abstract

The cytokine Interleukin 1 (IL-1) is an evolutionary innovation of vertebrates. Fish and amphibia have one *IL1* gene, while mammals have two copies of *IL1*, *IL1A* and *IL1B*, with distinct expression patterns and differences in their proteolytic activation. Our current understanding of the evolutionary history of IL-1 is mainly based on phylogenetic analyses, but this approach provides no information on potentially different functions of IL-1 homologs, and it remains unclear which biological activities identified for IL-1α and IL-1β in mammals are present in lower vertebrates. Here, we use *in vitro* and *in vivo* experimental models to examine the expression patterns and cleavage of IL-1 proteins from various species. We found that IL-1 in the teleost medaka shares the transcriptional patterns of mammalian IL-1α, and its processing also resembles that of mammalian IL-1α, which is sensitive to cysteine protease inhibitors specific for the calpain and cathepsin families. By contrast, IL-1 proteins in reptiles also include biological properties of IL-1β. Therefore, we propose that duplication of the ancestral IL1 gene led to segregation of expression patterns and protein processing that characterizes the two extant forms of IL-1 in mammals.

## 1. Introduction

The interleukin-1 (IL-1) family of cytokines orchestrates the immune response by mediating intercellular communication between many different cell types. Activated IL-1 has a range of inflammatory effects from fever induction to hematopoiesis and antibody synthesis (summarized in Dinarello, 2009). Like other immune-related cytokine genes, *IL1* genes are fast-evolving, driven by the need of the immune system to adapt to constantly changing threats. They have been identified in cartilaginous fish and all other jawed vertebrates (Bird et al., 2002; Rivers-Auty et al., 2018). Only one *IL1* gene is found in the genomes of most anamniotes (fishes and amphibians), although some teleost species such as rainbow trout (Heston, 1982) and carp (Engelsma et al., 2003) have two copies, most likely due to species-specific gene duplication events. The presence of *IL1A* and *IL1B* genes in all mammals and their localization on the same chromosome suggest that a tandem gene duplication event has occurred in their common ancestor (Eisenberg et al., 1991; Young and Sylvester, 1989).

The biological activities of IL-1α and IL-1β have been extensively analyzed in mice and humans. The two cytokines share a common transduction pathway but differ in their expression patterns and activation processes (Di Paolo and Shayakhmetov, 2016). At the transcriptional level, *IL1A* is constitutively expressed in a variety of cell types of hematopoietic and non-hematopoietic origin, such as keratinocytes, endothelial cells and the mucosal epithelium (Netea et al., 2015; Rider et al., 2017), whereas *IL1B* expression is predominantly induced in hematopoietic cells in response to inflammation (Dinarello, 2009). *IL1B* is also strongly expressed in various cancer cell types (Rébé and Ghiringhelli, 2020). The IL-1α protein is biologically active both in its full-length and cleaved forms, while the IL-1β full-length protein needs to be enzymatically cleaved to become active. The processing of the two IL-1 paralogs is regulated by distinct mechanisms. Both IL-1α and IL-1β can be processed by multiple proteases (Afonina et al., 2015). However, IL-1β is processed most efficiently by Caspase-1 (Thornberry et al., 1992), which, after its activation by the inflammasome (Martinon et al., 2002), cleaves IL-1β at two distinct sites (Howard et al., 1991). Caspase 1-mediated processing also results in the most bioactive form of IL-1β (Afonina et al., 2015). By contrast, Caspase-1 cannot process the IL-1α protein (Howard et al., 1991), which can instead be cleaved by Calpain proteases (Carruth et al., 1991; Kobayashi et al., 1990) and Granzyme B (Afonina et al., 2011). To what extent the biological activities of mammalian IL-1α and IL-1β are conserved in anamniotes is not known.

Thus far, the single *IL1* gene found in the genomes of lower vertebrates has been interpreted as being most closely related to mammalian *IL1B* and is therefore seen as a functional homolog. This assumption is mainly based on phylogenetic analyses (Ogryzko et al., 2014a; Rivers-Auty et al., 2018). However, the overall low conservation of IL-1 proteins between species justifies a reassessment of this interpretation and the consideration of other characteristic factors, such as gene expression patterns and protein processing mechanisms, to support a definite assignment. Here, we compare characteristics other than peptide sequences between IL-1 proteins of anamniotes and mouse IL-1α and IL-1β. We have created a reporter for *in vivo* visualization of the expression patterns and processing of IL-1 in transgenic medaka (*Oryzias latipes*) and tested *in vitro* the dependence of cleavage of IL-1 proteins from various anamniote species on Caspase-1. Our results show that the medaka ortholog is expressed and processed in a manner similar to mammalian IL-1α and that a conserved Caspase-1 cleavage site is already present in amniotes.

## 2. Results and Discussion

### 2.1. Evolution of IL-1 in vertebrates

Comparing nucleotide or amino acid sequences between species is a common method to elucidate evolutionary relationships. However, comparison of fast evolving genes across longer evolutionary times can be difficult, especially if pressures to diversify are active. In a phylogenetic tree of IL-1 proteins from lower and higher vertebrates, teleost IL-1 proteins form a separate cluster and share a branch point with clusters for mammalian IL-1α and IL-1β (Figure 1A), indicating that the amino acid sequences of IL-1α and IL-1β are equally distant from teleost IL-1, mostly consistent with what has been shown by other studies (Bird et al.; Gibson et al., 2014; Ogryzko et al., 2014a; Rivers-Auty et al., 2018). This is also true for avian and amphibian IL-1 proteins, which together form a separate cluster. Therefore, an accurate assignment of IL-1 proteins in lower vertebrates as homologs to either IL-1α or IL-1β on the basis of phylogenetic analysis is not possible. Another criterion that can help assign ancestral relationships of genes is the comparison of their genomic localization, i.e., synteny of neighboring genes across longer genomic stretches. The regions of vertebrate genomes in which the *IL1* genes are located are overall highly conserved, but this provides no helpful information because mammalian IL-1α and IL-1β are located next to each other within the same synteny group due to a tandem duplication event (Rivers-Auty et al., 2018). We therefore examined the conservation of characteristic amino acid sequences for IL-1α or IL-1β proteins that are relevant for their proteolytic processing. Alignment of IL-1 proteins from mammals, amphibians, reptiles, birds, teleosts, and cartilaginous fishes showed that known cleavage sites in mammalian IL-1α and IL-1β are poorly conserved in lower vertebrates (Figure 1B). Although all IL-1 proteins have the same structure in which the N- and the C-terminal domains are separated by a linker that contains potential cleavage sites (Figure 1C), many lower vertebrates lack the conserved aspartic acid residue as a Caspase-1 cleavage sites in this linker as well as the conserved beta trefoil fold that is characteristic for mammalian IL-1β (Figure 1B). The aspartic acid residue that at which the mammalian IL-1β N-terminus is cleaved is evolutionarily first present in the reptilian IL-1 protein (Supplementary Figure 1). Previous studies showed that the zebrafish IL-1 protein can be cleaved by Caspase A and Caspase B in transfected HEK cells (Li et al., 2018; Li et al., 2020; Vojtech et al., 2012). However, only one of the three aspartic acid residues identified as potential substrates of Caspase-1 homologues (Caspa and Caspb) in zebrafish IL-1 (Ogryzko et al., 2014a; Vojtech et al., 2012) can be cleaved by Caspase-1 in the sea bass (Reis et al., 2012). This aspartic acid residue in zebrafish IL-1 can be found in avian IL-1 but not in mammalian IL-1β. Besides the Caspase-1 site, the potential cleavage sites for other proteases such as Calpains, Cathepsins or Elastase are poorly conserved in IL-1 homologs (Figure 1B). Therefore, protein alignments are not sufficient to assign IL-1 genes in lower vertebrates as direct ancestors of either IL-1α or IL-1β in mammals or even to deduce the function of an IL-1 common ancestor. Also, to what extent the biological activity of IL-1 proteins in anamniotes depends on their processing is still unknown.

**Figure 1.**
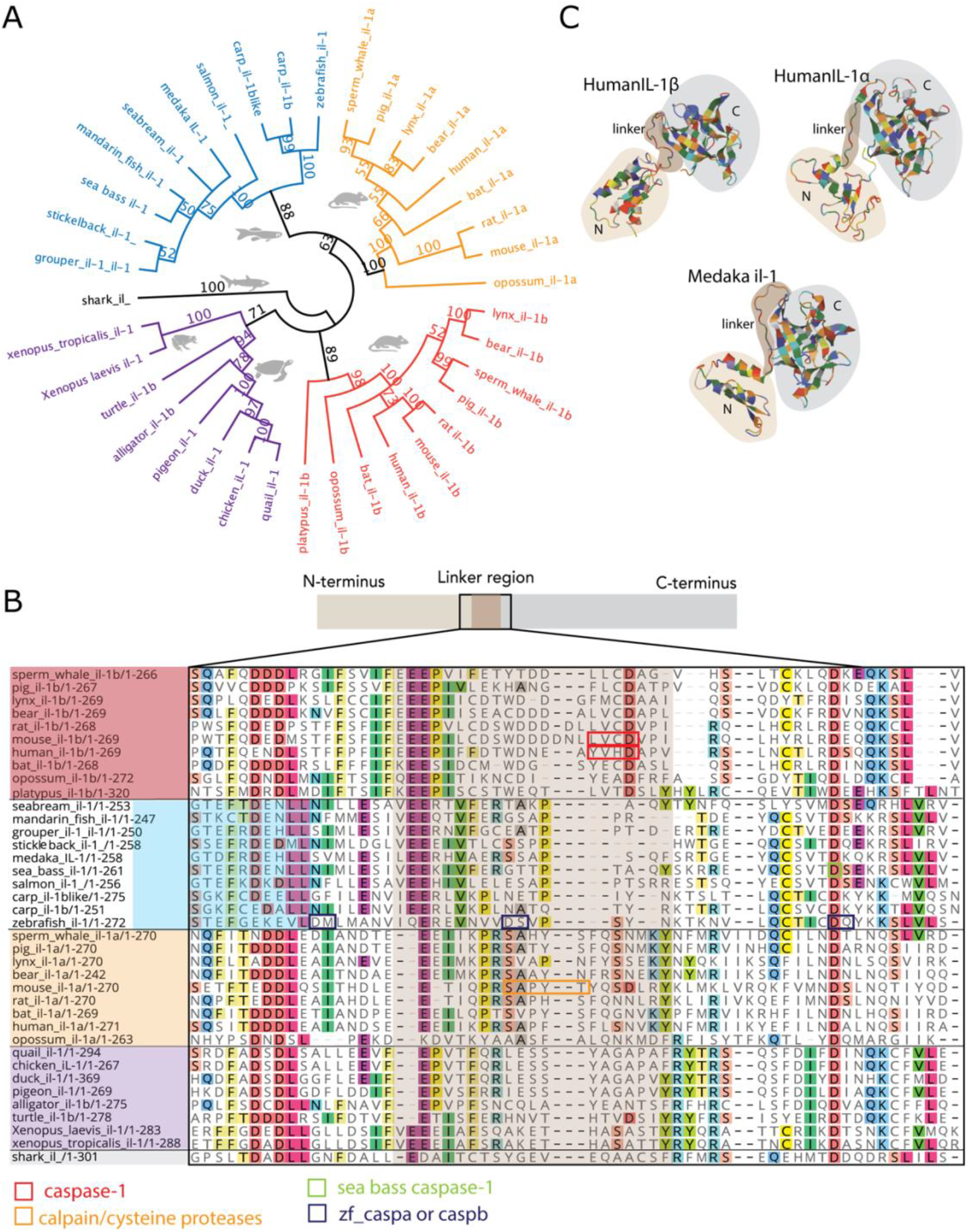
Phylogenetic analysis of IL-1 in vertebrates. (**A**) An unrooted tree calculated by Neighbor-Joining obtained from a Clustal W alignment of IL-1 full-length proteins. Calculated distance values are indicated for each branch. The accession numbers of genes used in this analysis are listed in the Supplementary Table 1. (**B**) An alignment of full-length IL-1 amino acid sequences from 28 species, showing the linker region (brown; corresponding to amino acids of linker as predicted by 3D structure of human IL-1β) and surrounding sequences. Experimentally confirmed IL-1 cleavage sites are marked with boxes as indicated. (**C**) Three-dimensional structures of medaka IL-1 compared to human IL-1α and IL-1β as predicted by RaptorX.

### 2.2. Expression of medaka il1 in naïve and upon infection or local injury

To better understand the evolutionary history of IL-1, we performed a comprehensive comparative analysis. One aspect that distinguishes mammalian *IL1A* and *IL1B* is their distinct expression profiles. *IL1A* is constitutively expressed at high level in various cell types, including epithelial and hematopoietic cells, while *IL1B* expression is weak but strongly inducible in monocytic cells in response to inflammation (Dinarello, 2009; Hadadi et al., 2016). To determine the expression activity of the *il1* gene in lower vertebrates, we use medaka as a model. We performed whole-mount *in situ* hybridization (WISH) with a probe for the *il1* full-length transcript but could not detect expression in naïve embryos. However, in embryos injected with *E. coli*, *il1* was strongly expressed around the site of injection (Supplementary Figure 2), which is consistent with previous observations in zebrafish (Ogryzko et al., 2014b). Because it was not clear whether absence of *il1* staining in naïve embryos is due to insufficient sensitivity of the WISH, we created an *il1* transgenic reporter fish, in which a 6.9 kb long *il1* promoter drives the transcription of a t2a-based bi-cistronic mRNA (Szymczak et al., 2004) encoding GFP and medaka full-length *il-1* tagged with a hemagglutinin (HA) peptide at the C-terminus (Figure 2A). This reporter allowed us not only to reveal the spatial expression patterns of *il1* gene but also to assess the processing of the IL-1 protein under various conditions using the HA-specific antibody. The GFP signal was detectable as a weak fluorescence signal in the epidermis of live embryos one day post-fertilization (dpf) (Figure 2B). At later stages, the GFP signal became restricted to the epithelial compartment of the skin, gills, and thymus as well as the neuromasts of the lateral line (Figure 2C). Similar to our observation, zebrafish *il1* is expressed in the skin, gills, and thymus (Hasegawa et al., 2017; Nguyen-Chi et al., 2014). Furthermore, human *IL1A* is expressed in keratinocytes and thymic epithelial cells (Dalloul et al., 1991). These findings suggest that *il1* expression in the epithelial compartment, which is characteristic for mammalian *IL1A*, is conserved among vertebrates.

**Figure 2.**
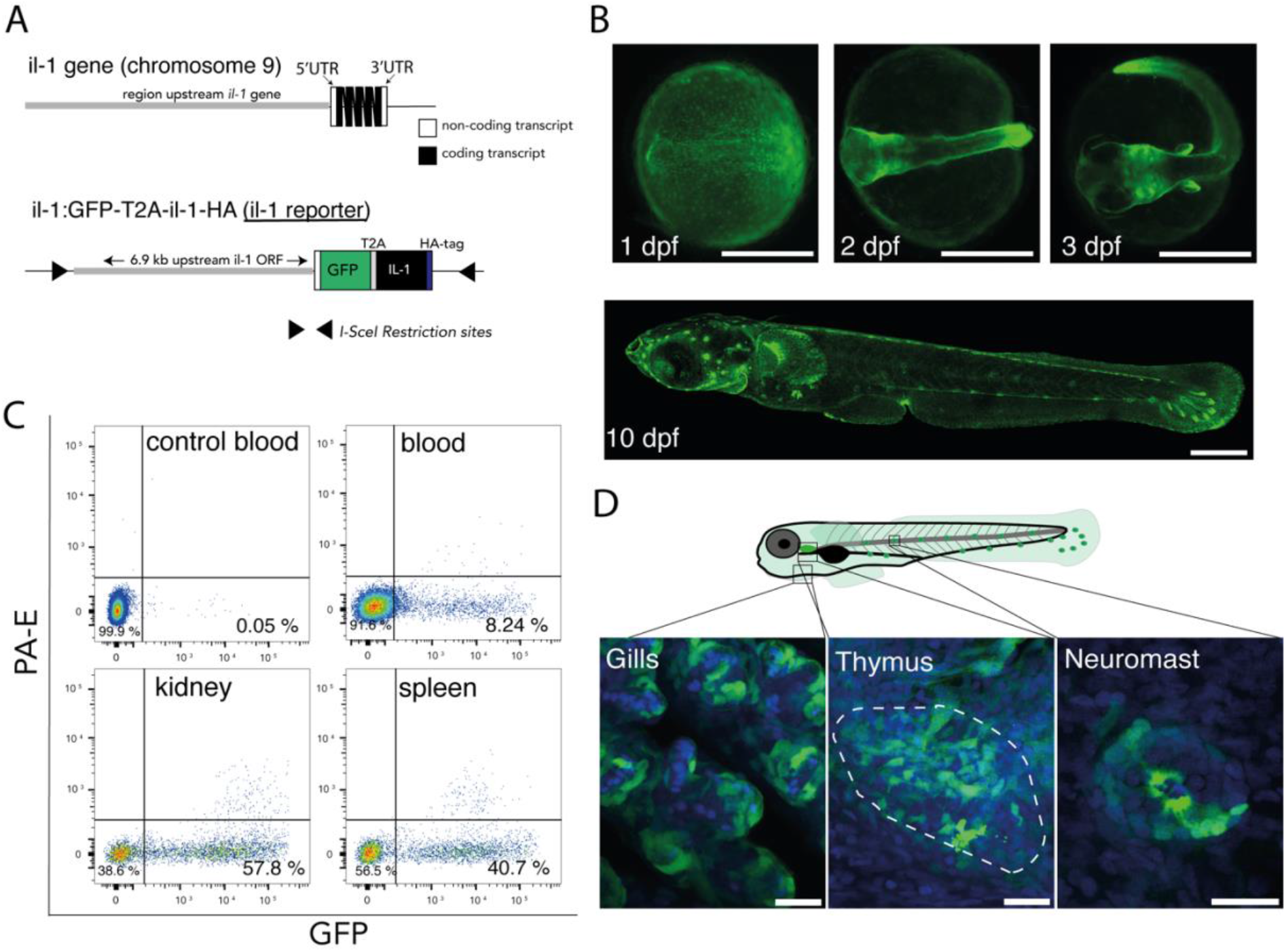
An *in vivo* reporter for medaka IL-1. (**A**) Top, schematic of the *il1* locus on medaka chromosome 9. Bottom, the transgenic il1 reporter construct indicating the position of 6.9 kb genomic fragments upstream of the *il1* gene that drives GFP and medaka full-length *il1* cDNA with a C-terminal HA tag. (**B**) GFP expression in the transgenic il1 reporter during embryogenesis. (**C**) Flow cytometry of hematopoietic cells isolated from blood, kidney, and spleen of naive adult il1 reporter fish. Blood from non-transgenic fish was used as a control. Data are representative of two independent biological samples. (**D**) GFP expression in the epithelial compartment of the gills, thymus and neuromast in the transgenic il1 reporter larvae. Nuclei are stained with DAPI (blue). Scale bars in B and D indicate 500 and 20 μm, respectively.

We also analyzed the *il1* expression in the adult hematopoietic cells and performed flow cytometry of isolated cells from blood, kidney, and spleen (Figure 2D). As negative control, blood of non-transgenic fish was used. Eight percent of blood cells, 57.8% of kidney cells, and 40.7% of spleen cells were GFP-positive. The constitutive expression of medaka *il1* in hematopoietic cells is consistent with zebrafish *il1* (Hasegawa et al., 2017; Nguyen-Chi et al., 2014) and mouse *IL1a* (Takacs et al., 1988). This result further supports the notion that regulatory elements controlling the constitutive expression of *IL1A* are also conserved in lower vertebrates.

Besides their constitutive expression, *il1* genes in lower vertebrates are inducible by inflammatory stimuli (Hasegawa et al., 2017; Nguyen-Chi et al., 2014). Our WISH analysis (Supplementary Figure 2) further confirms this observation. To distinguish whether *il1* inducibility is restricted to epithelial compartments or hematopoietic cells, we performed local injury and subcutaneous injection of bacteria in the transgenic fish. The GFP signal increased substantially in the epidermis when 50 μM Lipopolysaccharide (LPS) was injected into the muscle tissue (Figure 3A) or when the tail fins of transgenic larvae were injured (Figure 3B, C) indicating that *il1* expression can be induced in non-hematopoietic cells. Next, we subcutaneously injected bacterial debris conjugated with Alexa Fluor 594 into adult transgenic fish and analyzed hematopoietic cells, isolated from blood, kidney, and spleen using flow cytometry 16 hours-post-injection. We identified cells that expressed *il1* and had engulfed bioparticles by their combined red and green fluorescence (6.5%, 47.7% and 39.1% GFP^+^/RFP^+^ cells in blood, kidney, and spleen, respectively; data from two independent experiments). The presence of GFP^+^/RFP^+^ cells in the spleen (Figure 3D) indicated that that all myeloid cells that had engulfed bioparticles also expressed the *il1* gene. Whether *il1* expression was induced in them locally and they then migrated into the spleen, as a secondary lymphoid organ, to initiate the adaptive immune response cannot be deduced from this data. Taken together, our results reveal that *il1* is constitutively expressed in various epithelial tissues and can be upregulated in keratinocytes and myeloid cells upon infection or local injury. Therefore, the expression pattern of medaka *il1* resembles that of mammalian *IL1A* which is both constitutive and inducible (Chan et al., 2017; Luan et al., 2017; Suzuki et al., 2000).

**Figure 3.**
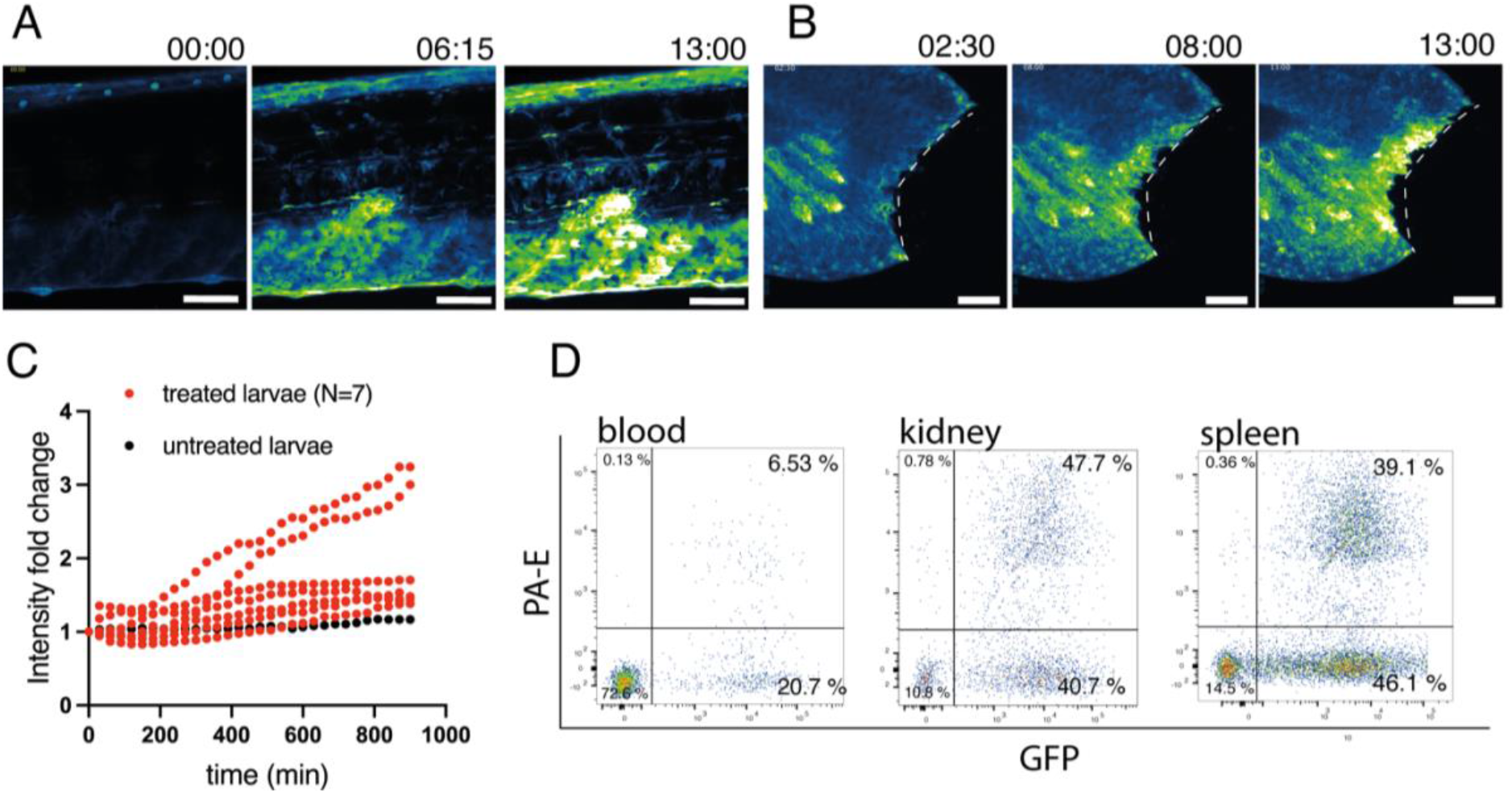
Induction of medaka *il1* upon injury and infection. (A) Still photographs from a time-lapse recording illustrating GFP upregulation in response to injection of 50 μM LPS into the muscle tissue of a transgenic il1 reporter larva. Numbers indicate time in hours. (**B**) Still photographs from a time-lapse recording illustrating GFP upregulation in response to a tail-fin cut of a transgenic il1 reporter larva. The dashed lines indicate the cut site. Numbers indicate time in hours. Scale bars in A and B indicate 50 μm. (**C**) The fold-change of mean GFP intensity quantified in the tail fin upon injury compared to untreated larvae. (**D**) Flow cytometry of hematopoietic cells isolated from blood, kidney, and spleen of il1 reporter fish 16 hours after subcutaneous injection of bacterial debris conjugated with Alexa 594. The data from Figure 2C and 3D come from the same experiment and the untreated group shown in Figure 2B is therefore the control for this panel. Data are representative of two independent experiments.

### 2.3. Processing of medaka IL-1 by proteases *in vivo*

To investigate the processing of medaka IL-1 in response to inflammatory stimuli, we used an anti-HA antibody to detect the transgenic, C-terminally tagged IL-1 in whole fish lysates on Western blots (WB). To prime immune cells, freshly hatched transgenic larvae were first exposed to LPS for 150 minutes (Figure 4). In a second treatment, we added either nigericin or ionomycin for an additional 45 minutes before lysates from whole larvae were prepared. Nigericin acts as a potassium ionophore that activates the NLRP3 inflammasome (Mariathasan et al., 2006), which is required for the Caspase-1-dependent cleavage and secretion of mammalian IL-1β (reviewed in: Franchi et al., 2009). By contrast, ionomycin is a membrane permeable calcium ionophore that increases intracellular calcium levels triggering Calpain activation and mature IL-1α release (Groβ et al., 2012; Tapia et al., 2019). Untreated larvae were used as a control group. The IL-1 pro-peptide is estimated to be around 29 kDA, and the C-terminal cleavage products are expected to be between 16.8 and 19.8 kDa if the precursor is cleaved within the linker region between the N- and C-terminal domains (the predicted products are schematically depicted in Figure 4B). Western blot analysis showed several bands for IL-1 (Figure 4C). In the control group, we detected HA-positive proteins around 29 and 58 kDa. The latter product is probably a read-through of the GFP-T2A-Il1 open reading frame that occurs when T2A-induced cleavage is not efficient (Kim et al., 2011). An additional protein fragment with a size around 20 kDa was detected when larvae were treated with 20 mM ionomycin (Figure 4C). This peptide was not detected upon treatment with LPS alone or LPS with the potassium ionophore nigericin, a commonly used inflammasome activator.

**Figure 4.**
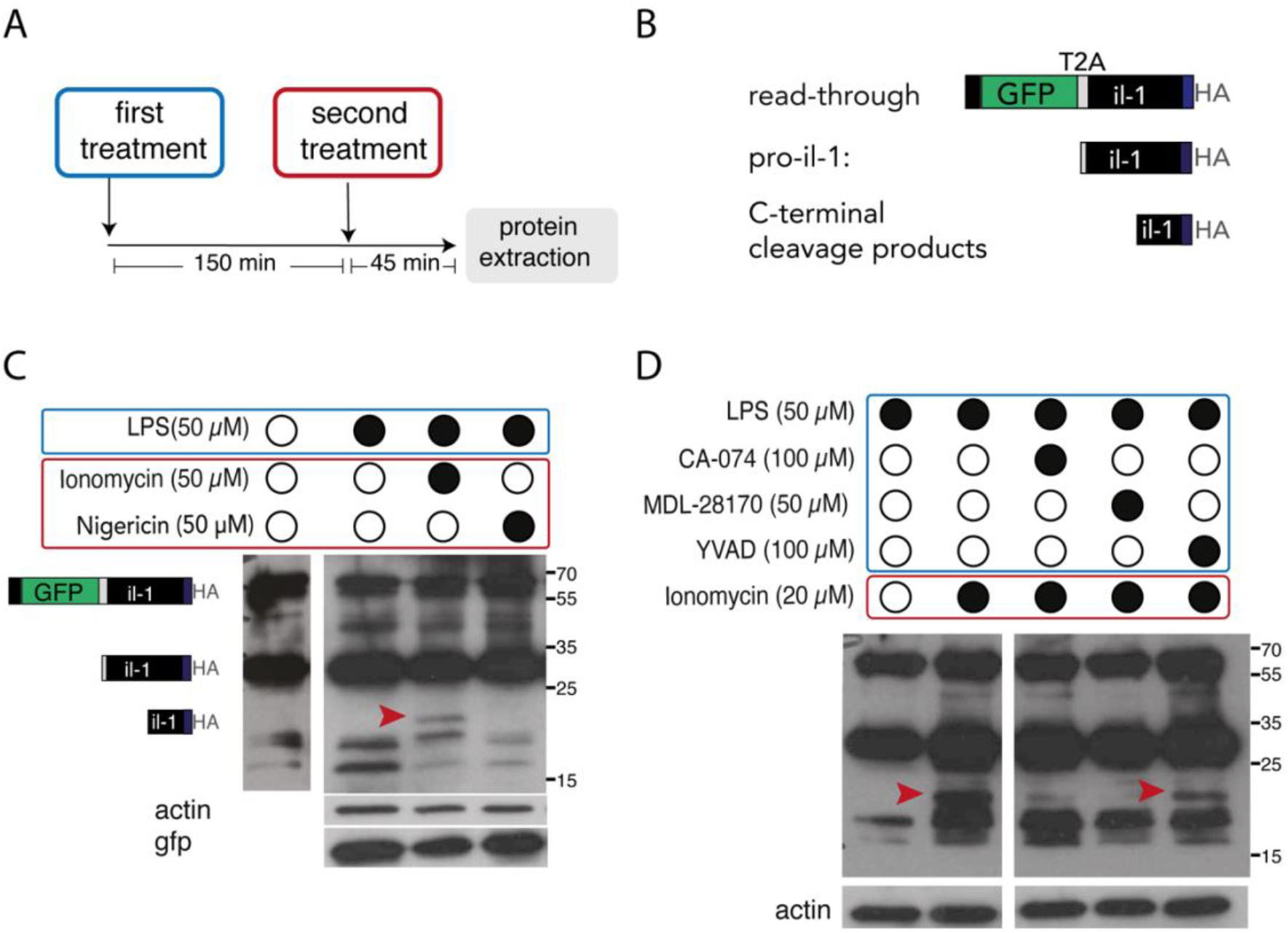
*In vivo* cleavage of medaka Il-1 upon chemical stimulation. (**A**) The experimental rationale. Freshly hatched medaka larvae were first treated with LPS for 150 minutes and then with a second compound (either nigericin or ionomycin) for an additional 30-45 minutes before protein extraction. (**B**) Schematic description of three predicted protein products that can be detected by HA antibody on Western blots. A fusion protein of GFP-t2a-*il1*-HA resulting from failure of the t2a induced ribosome skipping has a predicted MW of 58 kDa. The size of the Il-1 pro-peptide is 29 kDa. The molecular weight of cleaved Il-1 products was predicated between 16.8 and 20.8 kDa for a cleavage site located within the linker region. (**C**, **D**) WB analysis from medaka larvae after treatment with chemical compounds using HA, GFP and actin antibodies. Red arrowheads indicate the processed Il-1 protein. In (**D**), il1 reporter larvae were additionally treated with either cysteine protease inhibitors (MDL-28170 and CA-074) or caspase-1 inhibitor Y-VAD. Note the appearance of an Il-1 cleaved form only after LPS and Ionomycin treatment (**C**), which was reduced after MDL-28170 or CA-074, but not YVAD treatment (**D**). Data are representative of 4 independent experiments.

The 20 kDa IL-1 protein was still present when larvae were treated with the Caspase-1 inhibitor Ac-YVAD-cmk. By contrast, when larvae were treated with cysteine protease inhibitors MDL-28170 and CA-074 prior to and during ionomycin treatment, the 20 kDa protein was not detectable. Given that MDL-28170 and CA-074 inhibit Calpains and proteases of the Cathepsin family (Mehdi, 1991; Montaser et al., 2002), our result indicates that medaka IL-1 can be processed by cysteine proteases from one or both of these protein families.

We also assessed the spatial expression patterns of *calpains* and their small subunit *capns1* as well as *cathepsin B*, *L* and *S* in medaka embryos. WISH analysis showed that *calpain2*, *capns1b* and c*athepsin L2* and *S* are all expressed in the skin and gut, with enhanced expression in neuromasts (Supplementary Figure 4). Their colocalization with *il1* expression makes them potential candidates for IL-1 processing enzymes in medaka.

### 2.4. Processing of medaka IL-1 in vitro

To further test our conclusion that medaka IL-1 is processed in a similar manner as mammalian IL-1α, we compared the processing of medaka IL-1 and mouse IL-1α and IL-1β in a cell-based assay using the pro-interleukin-1b-Gaussia luciferase (iGLuc) fusion assay (Bartok et al., 2013). In this assay, pro-IL-1-dependent formation of protein aggregates renders the Gaussia luciferase (GLuc) inactive, and this can be reversed if the cytokine is cleaved, leading to recovery of luciferase activity (Figure 5A). We transfected mouse J774 macrophages with constructs containing full-length cDNAs of medaka *il1*, mouse *IL1a* or mouse *IL1b* fused with the Gluc reporter. Transfected macrophages were then treated in a similar protocol as in the *in vivo* experiments (Figure 4A). *In vitro*, LPS alone was not sufficient to induce luciferase activity in macrophages transfected with any of the three IL-1 constructs. However, luciferase became activated when transfected cells were treated with nigericin or ionomycin. Consistent with our previous study (Bartok et al., 2013), luciferase was activated up to 50-fold when cells transfected with mouse IL1b-Gluc were treated with nigericin (Figure 5B), but not with ionomycin. The effect of nigericin on cleavage of mouse IL-1α and medaka IL-1 was lower. Conversely, ionomycin treatment resulted in a strong luciferase activity in cells transfected with mouse IL-1α-Gluc or medaka IL-1-Gluc constructs (Figure 5B).

**Figure 5.**
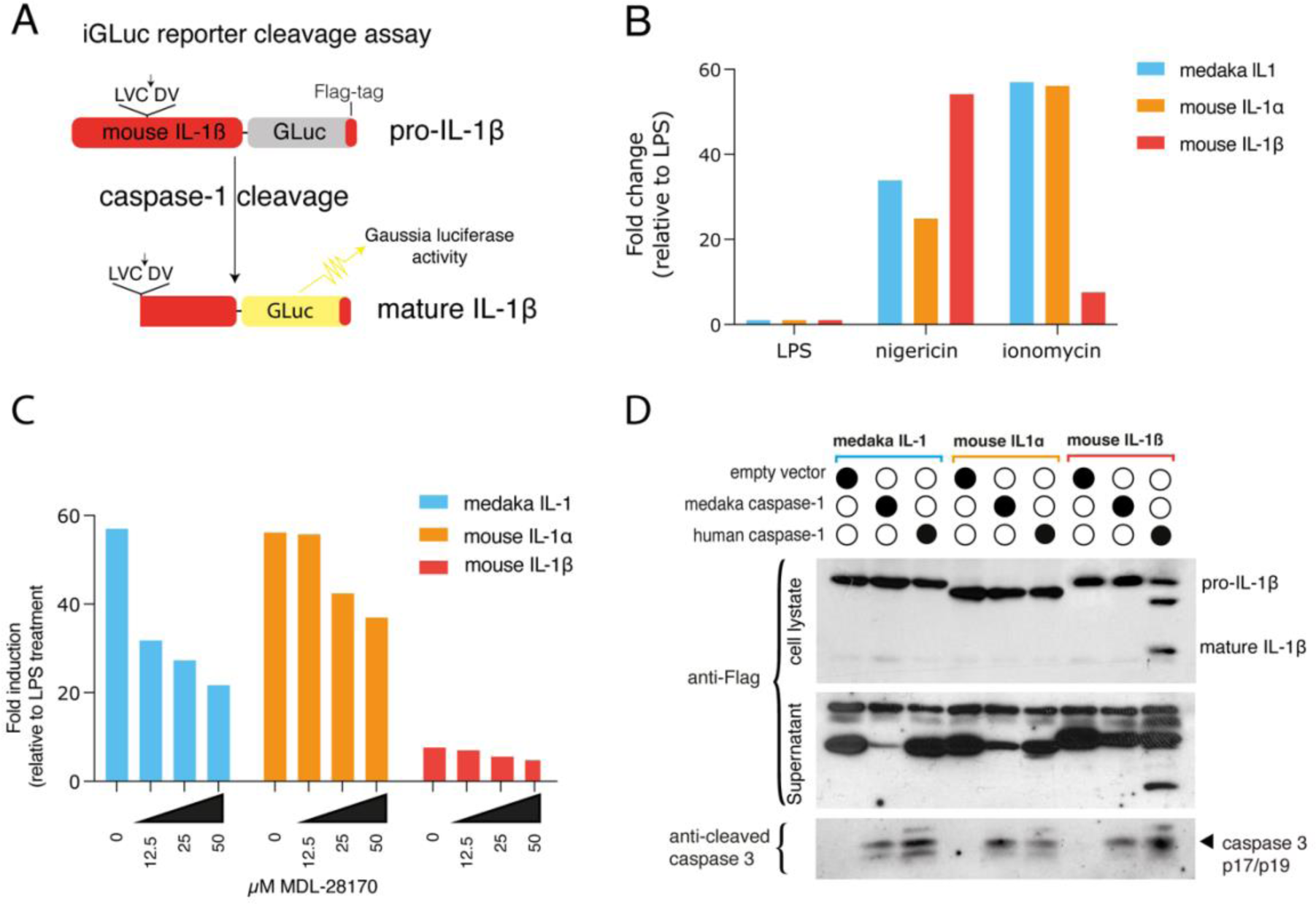
In vitro cleavage of mouse IL-1α, IL-1β and medaka Il-1 upon treatment. (**A**) Schematic description of the iGLuc reporter cleavage assay (Bartok et al., 2013). In this assay, the pro-IL-1β-GLuc fusion protein forms aggregates and is enzymatically inactive. The cleavage of pro-IL-1β-GLuc by Caspase-1 results in monomeric and enzymatically active protein. (**B**) Relative fold-change of luciferase activity upon treatment with different chemical compounds compared to cells treated with only LPS. Mouse J774 macrophages were transfected with pro-IL-1β-GLuc, pro-IL-1α-GLuc and medaka Il-1-GLuc constructs. Data are representative of 4 independent experiments. (**C**) Relative fold-change of luciferase activity in transfected J774 cells with different iGLuc constructs and subsequent treatment with ionomycin and MDL-28170. Transfected cells treated with only LPS were used as a control group. Data are representative of 4 independent experiments. (**D**) WB analysis of transfected HEK293T cells with different constructs expressing IL-1 and Caspase-1 cDNAs from the indicated species. Detection of cleaved endogenous Caspase-3 in transfected cells indicates autoactivation of Caspase-1 in 293T cells, which do not express GSDMD (Heilig et al., 2020; Masumoto et al., 2003; Tsuchiya et al., 2019).

Mouse IL-1α has been reported to be cleaved by cysteine proteases. To determine whether this is also true of medaka IL-1, we additionally applied the cysteine protease inhibitor MDL-28170 along with LPS and ionomycin. Here, we found a dosage-dependent decrease of luciferase activity for constructs carrying medaka IL-1 or mouse IL-1α (Figure 5C), suggesting that the cleavage of medaka IL-1 also depends on cysteine proteases and further confirming our *in vivo* observations. To test whether Caspase-1 can cleave medaka IL-1, we co-transfected 293T cells with plasmids carrying either medaka or human Caspase-1. We found that human Caspase-1 was only able to cleave mouse IL-1β but not mouse IL-1α or medaka IL-1 (Figure 5D). Medaka Caspase-1 was not able to cleave any of the tested IL-1 proteins. Together, these results indicate that medaka IL-1 and mouse IL-1α can be processed by cysteine proteases of the calpain or cathepsin family.

### 2.5. Caspase-1 mediated IL-1 cleavage in vertebrates

The processing of IL-1β by Caspase-1 seen in mammals does not appear to occur in medaka IL-1. However, zebrafish IL-1 can be processed by zebrafish Caspase A (also named Casp1) and Caspase B (also named Casp19a) in transfected HEK cells (Li et al., 2018; Li et al., 2020; Vojtech et al., 2012) and primary zebrafish leukocytes (Vojtech et al., 2012). It is worth nothing that the zebrafish inflammatory Caspases (Caspase A, Caspase b, Caspase 19b and Caspase 23) differ from mammalian Caspase-1 and the Caspase-1 found in other teleost species: while the latter have a Caspase recruitment domain (CARD) at their N-terminus, zebrafish Caspases A, B and 19b have a Pyrin (PYD) domain instead. Moreover, the mutually dependent activity of Caspase A and B necessary for cleavage of zebrafish IL-1 is not conserved in other vertebrates, and the aspartic acid residues identified by (Vojtech et al., 2012) as Caspase-A and Caspase-B-specific cleavage sites are not conserved Caspase-1 cleavage sites in mammals.

Therefore, it is likely that zebrafish has independently acquired the ability to be cleaved by caspases. The alignment of IL-1 proteins in vertebrates shows that the N-terminal mammalian IL-1β Caspase-1 cleavage site is conserved in amniotes (Figure 6A). To determine whether Caspase-1-mediated IL-1 cleavage is characteristic for amniotes, we tested IL-1 proteins from different amniotes (reptiles: alligator and turtle) and anamniotes (fish: shark; amphibian: Xenopus). By co-transfecting IL-1 constructs with human Caspase-1, we found that both turtle and alligator IL-1 are cleaved at a site close to the N-terminus, estimated by the product size of around 30 kDa, similar to the intermediate cleavage product of mouse IL-1β (Figure 6B). By contrast, IL-1 of Xenopus and shark could not be cleaved by human Caspase-1. Taken together we show that, first, the expression patterns and protein cleavage of IL-1 in medaka resemble the mammalian IL-1α and, secondly, the cleavage of IL-1 by Caspase-1 observed has evolved in amniotes. Therefore, a designation of IL-1 in anamniotes as homologue of IL-1β is currently not justified. Additional experimental models will be needed to elucidate the extent to which the cleavage of IL-1 proteins by calpains in anamniotes is necessary for their activity.

**Figure 6.**
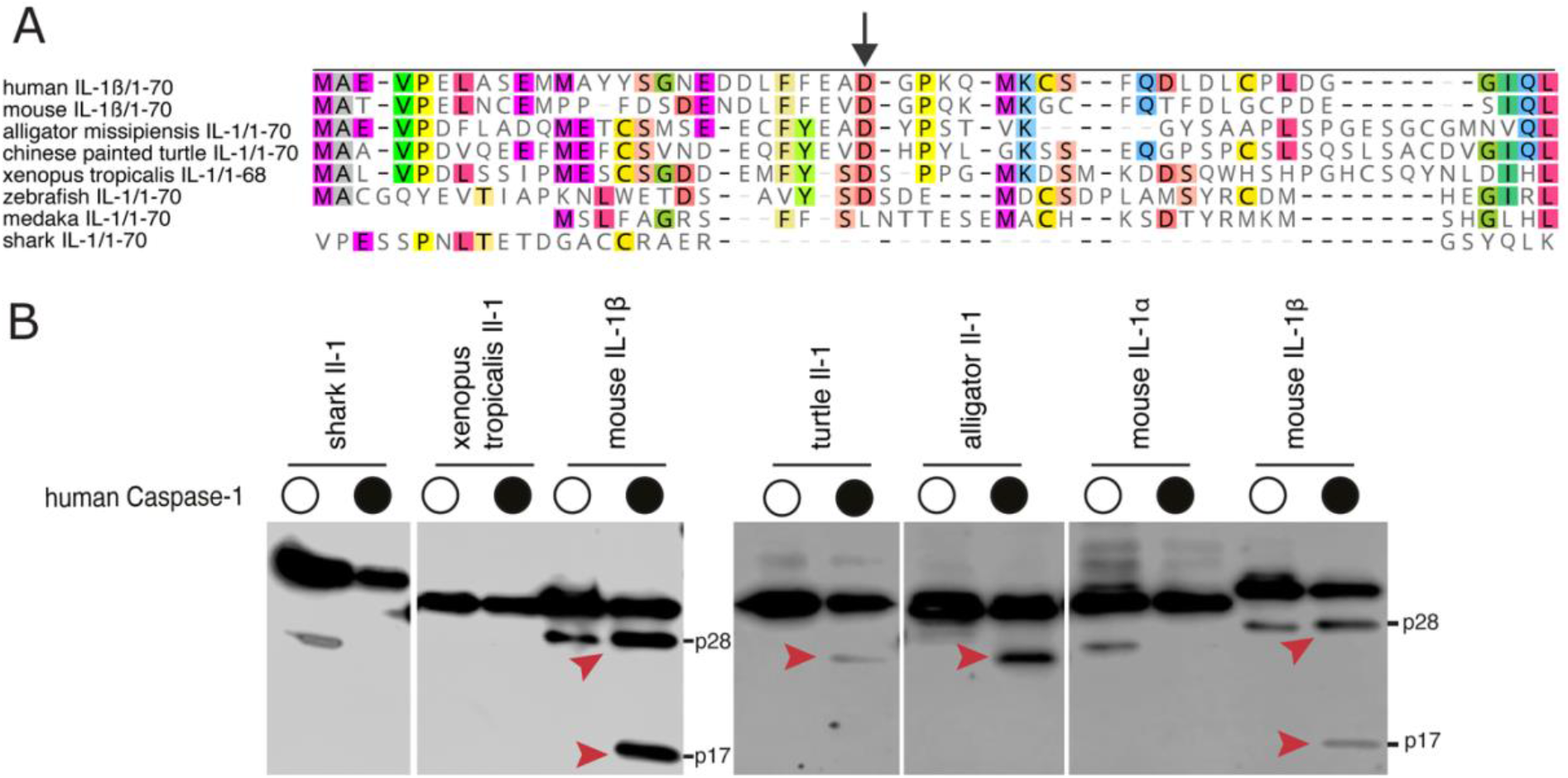
Caspase-1 dependent cleavage of IL-1 in amniotes. (**A**) Alignment of the first 70 amino acids of amniote (human, mouse) and anamniote species, showing a conserved aspartic acid residue (indicated by arrow) between mammalian IL-1β and reptiles IL-1. The accession numbers of genes used are listed in the Supplementary Table 1. (**B**) *In vitro* Caspase-1 assay showing cleavage of turtle and alligator IL-1 by human Caspase-1. Red arrowheads indicate Caspase-1-specific cleavage products.

## 3. Material and Methods

### 3.1. Bioinformatic analysis

Sequences were retrieved using BLASTP searches (http:/www.ncbi.nlm.nih.gov/ or ensemble.org) with default parameters using human and mouse IL-1 proteins. In our phylogenetic tree analysis, we included IL-1 protein sequences from nine mammals, two reptiles, four birds, eight teleosts and one cartilaginous fish. All genes are listed in Supplementary Table 1. Sequence alignment and phylogenetic trees were done with the Geneious (version 3) software.

### 3.2. Fish

Adult medaka (*Oryzias latipes*) were kept under a 14 h light – 10 h dark cycle at 26°C. Embryos were collected and kept in embryonic rearing medium (ERM). Freshly hatched yolk-sac transgenic larvae were used for most of three experiments. Generation of medaka transgenic reporter lines, and all experimental protocols were performed in accordance with relevant institutional and national guidelines and regulations and were approved by the EMBL Institutional Animal Care and Use Committee (IACUC nos. 2019-03-19ML).

### 3.3. Generation of transgenic fish

To generate transgenic il1:gfp-t2a-*il1*-HA reporter fish, a fragment containing GFP and full-length of medaka il1 cDNA separated by t2a, a short viral sequences, were cloned into a vector containing 6.9 kb upstream region of the *il1* gene (Figure 2A). The plasmid at 10-25 ng/μl concentration together with 1 μl *I-SceI* meganuclease and NEB buffer (NewEngland BioLabs) were co-injected into the blastomere at one-cell stage embryos. F0 larvae with GFP signal were selected for breeding.

### 3.4. Immunohistochemistry

Larvae were fixed with 4 % paraformaldehyde (PFA) in 0.1% Tween PBS (1xPTW). After three washes, larvae were incubated in 30% sucrose/PTW for 24 hrs followed by 50% tissue freezing medium/30%Sucrose/PTW for another day. Samples were mounted and sectioned at 20 μM on a cryostat (Leica Biosystems CM2050S). Sections were rehydrated for 20 min with 1x PTW and blocked with 10% NGS/PTW for 2 hrs. They were incubated with 1:500 mouse-anti-GFP (Sigma) and 1:500 Rb-anti-HA antibody (Cell signaling) in 1% NGS/PTW over night at 4°C. After several washing steps in PTW, sections were stained with anti-Rabbit-Alexa 647 and anti-mouse-Alexa488 in 1% NGS/PTW with 1:1000 diluted DAPI for 2 hrs at 37°C. Slides were washed 3x 10 min with PTW and then mounted with Vectra shield (Vectra labs). A Zeiss 780 confocal microscope with a 40x water objective was used for imaging of stained sections.

### 3.5. Flow cytometry

Cells were isolated from spleen, kidney, and blood from adult transgenic fish. To avoid blood coagulation, ice cold 0.57x PBS with 30 mM EDTA was used to collect blood from fish. Cells were disaggregated using a cell strainer (40 μm Nylon, BD Falcon) and collected in FACS buffer (5 mM EDTA, 10 U/ml Heparin in 1x PBS). The BD LSR Fortessa Cell Analyzer (BD Biosciences) was used for flow cytometry analysis.

### 3.6. WISH

Whole-mount RNA *in situ* hybridization (WISH) was performed as described previously (Aghaallaei et al., 2007) Probes used in this study are listed in Supplementary Table 2.

### 3.7. Wounding assay

The wounding assay was adapted from de Oliveira et al. 2013. Briefly, freshly hatched yolksac larvae were anaesthetized in 40 μg/ml ethyl-m-aminobenzoate methansulfonate (tricaine) in ERM. The caudal fin of larvae was cut using sterile surgical blades. Larvae were then immediately mounted in 1% low-melting agarose containing 40 μg/ml tricaine and live imaged overnight using confocal microscopes (Zeiss 780 NLO or Leica SP8). The fluorescence intensity over time was calculated using SUM intensity projections, background subtraction, intensity was measured along the line of the cut side or along the rim of the fin in uninjured fins. After background subtraction, the fold change was calculated from signal intensity at t_x_ divided by initial (t_0_) fluorescence intensity (t=time).

### 3.8. Injection of LPS and bacteria

Anesthetized larvae were subcutaneously injected with 50 μg/ml Lipopolysaccaride (LPS, Sigma) using a glass needle. Anesthetized adults were injected with PBS containing *Staphylococcus aureus* BioParticles™ Alexa Fluor 594 conjugate (ThermoFisher). No fish died as a result of the injection. Adult fish were then kept separately in tanks for 16 hours before euthanization and sample preparation for flow cytometry.

### 3.9. *In vivo* IL-1 cleavage assay

Freshly hatched yolksac larvae were incubated in 3-4 ml ERM substituted with different combinations of compounds in a 6-well plate as shown in Figure 4A. Larvae were first treated with 50 μg/ml LPS for 2,5 hours followed by 20-50 μm ionomycin (Cayman Chemicals) or 50 μm nigericin (Sigma) for additional 20-60 min. The treatment was terminated when larvae showed clear signs of exposure (immobility). Inhibitors MDL-28170 (Santa Cruz), CA-74 (Cayman Chemicals) and Ac-YVAD-cmk (Sigma) were added directly to the LPS containing medium and the concentration was kept constant after adding ionomycin into the medium. After treatment, each larva was transferred into a 1.5 ml tube on ice for subsequent protein extraction. 30 μl protein extraction buffer (10 mM HEPES, 100 mM KCl, 2 mM MgCl_2_, 0,1 mM CaCl_2_, 5 mM EGTA, pH=8.0, 0.9 mM TritonX ,1 mM NaF, 1 mM Na_3_VO_4_ and proteinase inhibitor) was then added to the tube. Samples were squished using a pestle. The suspension was kept on ice for 20 min and then centrifuged for 20 min 4°C at 10.000 rpm. The supernatant was transferred into a new tube and stored at −20 °C. To detect proteins, heat denatured larval suspension were run on a 15% SDS-PAGE and then transferred into a 0.45 μM nitrocellulose membrane by semi-dry electroblotting for 35 min at 13 V. Blots were incubated with 1:1000 anti-HA antibody (Rabbit, Cell signaling), 1:20000 anti-GFP (mouse, Sigma) and 1:20,000 anti-actin (rabbit, Sigma).

### 3.10. *In vitro* IL-1 cleavage assay

For lentiviral overexpression in J774 macrophages, murine IL1a-GLuc, murine IL1b Gluc and medaka IL1 were subcloned into 3rd generation lentivector pLenti6-EF1alpha-IRES-EGFP (a derivative of Invitrogen pLenti6, kindly provided by Jonas Doerr, Institute of Reconstructive Neurobiology, University of Bonn) via SalI/NotI fusion. Lentiviruses were generated using calcium-phosphate transfection of HEK293T, and J774 macrophages were spin transduced, as described in (Kutner et al., 2009) and sorted for GFP expression. For inflammasome experiments, the luciferase signal was measured directly from the supernatant after addition of the Gaussia luciferase substrate coelenterazine as performed in (Bartok et al., 2013).

For transient transfection, murine IL1a-GLuc, murine IL1b Gluc and medaka IL1 were subcloned into the mammalian expression vector pEFBOS containing a C-term FLAG-tag via *XhoI*/*BamHI* fusion. Medaka caspase-1 and human caspase-1 were subcloned into pLenti6-EF1alpha-IRES-EGFP. HEK293T cells were transfected with the indicated plasmids using TransIT-LT1 (Mirus Bio). Cells were lysed with SDS-Sample buffer 24h after transfection and prepared for immunoblotting. To detect proteins, heat denatured samples were run on a 12% SDS-PAGE and then transferred into a 0.2 μM nitrocellulose membrane using wet transfer (50min, 100V). Blots were incubated with 1:1000 Monoclonal ANTI-FLAG® M2-HRP antibody (mouse, Sigma) or 1:1000 anti-cleaved Caspase-3 #9661 (rabbit, Cell Signaling Technology), followed by 1:3000 goat anti-rabbit HRP 1706515 (BioRad).

### 3.11. Statistical analysis

Wilcoxon-Mann-Whitney test was used to calculate significant differences where indicated. A *p*-value < 0.05 was considered statistically significant. The numbers of biological samples (N) for experiments are indicated in each figure. Data in bar graphs are shown as an absolute number with means ± S.D. noted. All data was analyzed in GraphPad Prism software (version 9).

## Authors’ contribution

BB, EB and ML made initial observations and designed the work. EHC, EB and HS performed analysis. EHC, EB, BB, and ML analyzed the data, prepared the draft and final version of the manuscript. All authors reviewed the results and approved the manuscript.

## Acknowledgments

The authors thank the Advanced Light Microscopy Facility (AMLF) at the EMBL-Heidelberg for their continued support. The Flow Cytometry Core Facility of the Medical Faculty at the University of Bonn for providing support and instrumentation funded by the Deutsche Forschungsgemeinschaft (DFG, German Research Foundation) – Project number 387333827. We also thank Jonas Doerr for providing a modified pLenti-IRES-EGFP plasmid for experiments, Saskia Schmitz for expert technical assistance, and Christine Gottschalk for performing WISH analysis.

## References

Afonina, I. S., Tynan, G. A., Logue, S. E., Cullen, S. P., Bots, M., Lüthi, A. U., Reeves, E. P., McElvaney, N. G., Medema, J. P., Lavelle, E. C., et al. (2011). Granzyme B-dependent proteolysis acts as a switch to enhance the proinflammatory activity of IL-1α. Mol. Cell 44, 265–278.

Afonina, I. S., Müller, C., Martin, S. J. and Beyaert, R. (2015). Proteolytic Processing of Interleukin-1 Family Cytokines: Variations on a Common Theme. Immunity 42, 991–1004.

Aghaallaei, N., Bajoghli, B. and Czerny, T. (2007). Distinct roles of Fgf8, Foxi1, Dlx3b and Pax8/2 during otic vesicle induction and maintenance in medaka. Dev. Biol. 307, 408–420.

Bartok, E., Bauernfeind, F., Khaminets, M. G., Jakobs, C., Monks, B., Fitzgerald, K. A., Latz, E. and Hornung, V. (2013). iGLuc: a luciferase-based inflammasome and protease activity reporter. Nat Methods 10, 147–154.

Bird, S., Zou, J., Wang, T., Munday, B., Cunningham, C. and Secombes, C. J. Evolution of interleukin-1?? Cytokine Growth Factor Rev. 13, 483–502.

Carruth, L. M., Demczuk, S. and Mizel, S. B. (1991). Involvement of a calpain-like protease in the processing of the murine interleukin 1α precursor. J. Biol. Chem. 266, 12162–12167.

Chan, J., Maninjay, A., Jiang, Z., Carpenter, S., Aiello, D., Elling, R., Fitzgerald, K. A. and Caffrey, D. R. (2017). A Natural Antisense Transcript, AS-IL1α, controls inducible transcription of the pro-inflammatory cytokine IL-1α. jimmunol 195, 1359–1363.

Dalloul, A. H., Arock, M., Fourcade, C., Hatzfeld, A., Bertho, J. M., Debre, P. and Mossalayi, M. D. (1991). Human thymic epithelial cells produce interleukin-3. Blood 77, 69–74.

Di Paolo, N. C. and Shayakhmetov, D. M. (2016). Interleukin 1α and the inflammatory process. Nat. Immunol. 17, 906–913.

Dinarello, C. A. (2009). Immunological and Inflammatory Functions of the Interleukin-1 Family. Annu. Rev. Immunol. 27, 519–550.

Eisenberg, S. P., Brewer, M. T., Verderber, E., Heimdal, P., Brandhuber, B. J. and Thompson, R. C. (1991). Interleukin 1 receptor antagonist is a member of the interleukin 1 gene family: evolution of a cytokine control mechanism. Proc. Natl. Acad. Sci. U. S. A. 88, 5232–6.

Engelsma, M. Y., Stet, R. J., Saeij, J. P. M. and Verburg-Van Kemenade, B. M. L. (2003). Differential expression and haplotypic variation of two interleukin-1β genes in the common carp (Cyprinus carpio L.). Cytokine 22, 21–32.

Franchi, L., Eigenbrod, T., Muñoz-Planillo, R. and Nuñez, G. (2009). The inflammasome: A caspase-1-activation platform that regulates immune responses and disease pathogenesis. Nat. Immunol. 10, 241–247.

Gibson, M. S., Kaiser, P. and Fife, M. (2014). The chicken IL-1 family: Evolution in the context of the studied vertebrate lineage. Immunogenetics 66, 427–438.

Groß, O., Yazdi, A. S., Thomas, C. J., Masin, M., Heinz, L. X., Guarda, G., Quadroni, M., Drexler, S. K. and Tschopp, J. (2012). Inflammasome Activators Induce Interleukin-1α Secretion via Distinct Pathways with Differential Requirement for the Protease Function of Caspase-1. Immunity 36, 388–400.

Hadadi, E., Zhang, B., Baidzajevas, K., Yusof, N., Puan, K. J., Ong, S. M., Yeap, W. H., Rotzschke, O., Kiss-Toth, E., Wilson, H., et al. (2016). Differential IL-1β secretion by monocyte subsets is regulated by Hsp27 through modulating mRNA stability. Sci. Rep. 6, 1–13.

Hasegawa, T., Hall, C. J., Crosier, P. S., Abe, G., Kawakami, K., Kudo, A. and Kawakami, A. (2017). Transient inflammatory response mediated by interleukin-1β is required for proper regeneration in zebrafish fin fold. Elife 6, 1–22.

Heilig, R., Dilucca, M., Boucher, D., Chen, K. W., Hancz, D., Demarco, B., Shkarina, K. and Broz, P. (2020). Caspase-1 cleaves Bid to release mitochondrial SMAC and drive secondary necrosis in the absence of GSDMD. Life Sci. Alliance 3, 1–15.

Heston, W. E. (1982). Genetics: animal tumors. In Cancer: a Comprehensive Treatise. 2nd ed. (ed. Becker, F..) Plenum Press.

Howard, A. D., Kostura, M. J., Thornberry, N., Ding, G. J., Limjuco, G., Weidner, J., Salley, J. P., Hogquist, K. A., Chaplin, D. D. and Mumford, R. A. (1991). IL-1-converting enzyme requires aspartic acid residues for processing of the IL-1 beta precursor at two distinct sites and does not cleave 31-kDa IL-1 alpha. J. Immunol. 147, 2964–9.

Kim, J. H., Lee, S.-R., Li, L.-H., Park, H.-J., Park, J.-H., Lee, K. Y., Kim, M.-K., Shin, B. A. and Choi, S.-Y. (2011). High cleavage efficiency of a 2A peptide derived from porcine teschovirus-1 in human cell lines, zebrafish and mice. PLoS One 6, e18556.

Kobayashi, Y., Yamamoto, K., Saido, T., Kawasaki, H., Oppenheim, J. J. and Matsushima, K. (1990). Identification of calcium-activated neutral protease as a processing enzyme of human interleukin 1 alpha. Proc. Natl. Acad. Sci. U. S. A. 87, 5548–52.

Kutner, R. H., Zhang, X. Y. and Reiser, J. (2009). Production, concentration and titration of pseudotyped HIV-1-based lentiviral vectors. Nat. Protoc. 4, 495–505.

Li, J., Gao, K., Shao, T., Fan, D., Hu, C., Sun, C., Dong, W., Lin, A., Xiang, L. and Shao, J. (2018). Characterization of an NLRP1 Inflammasome from Zebrafish Reveals a Unique Sequential Activation Mechanism Underlying Inflammatory Caspases in Ancient Vertebrates. J. Immunol. ji1800498.

Li, J.-Y., Wang, Y.-Y., Shao, T., Fan, D.-D., Lin, A.-F., Xiang, L.-X. and Li-Xin Xiang, and J.-Z. S. (2020). The zebrafish NLRP3 inflammasome has functional roles in ASC-dependent interleukin-1 maturation and gasdermin E – mediated pyroptosis. JBC 295, 1120–1141.

Luan, L., Patil, N. K., Guo, Y., Hernandez, A., Bohannon, J. K., Fensterheim, B. A., Wang, J., Xu, Y., Enkhbaatar, P., Stark, R., et al. (2017). Comparative Transcriptome Profiles of Human Blood in Response to the Toll-like Receptor 4 Ligands Lipopolysaccharide and Monophosphoryl Lipid A. Sci. Rep. 7, 1–16.

Mariathasan, S., Weiss, D. S., Newton, K., McBride, J., O’Rourke, K., Roose-Girma, M., Lee, W. P., Weinrauch, Y., Monack, D. M. and Dixit, V. M. (2006). Cryopyrin activates the inflammasome in response to toxins and ATP. Nature 440, 228–232.

Martinon, F., Burns, K. and Tschopp, J. (2002). The Inflammasome: A molecular platform triggering activation of inflammatory caspases and processing of proIL-β. Mol. Cell 10, 417–426.

Masumoto, J., Zhou, W., Chen, F. F., Su, F., Kuwada, J. Y., Hidaka, E., Katsuyama, T., Sagara, J., Ngo-hazelett, P., Postlethwait, J. H., et al. (2003). Caspy, a Zebrafish Caspase, Activated by ASC Oligomerization Is Required for Pharyngeal Arch Development *. 278, 4268–4276.

Mehdi, S. (1991). Cell-penetrating inhibitors of calpain. Trends Biochem. Sci. 16, 150–153.

Montaser, M., Lalmanach, G. and Mach, L. (2002). CA-074, But Not Its Methyl Ester CA-074Me, Is a Selective Inhibitor of Cathepsin B within Living Cells. 383, 1305–1308.

Netea, M. G., van de Veerdonk, F. L., van der Meer, J. W. M., Dinarello, C. A. and Joosten, L. A. B. (2015). Inflammasome-Independent Regulation of IL-1-Family Cytokines. Annu. Rev. Immunol. 33, 49–77.

Nguyen-Chi, M., Phan, Q. T., Gonzalez, C., Dubremetz, J. F., Levraud, J. P. and Lutfalla, G. (2014). Transient infection of the zebrafish notochord with E. coli induces chronic inflammation. DMM Dis. Model. Mech. 7, 871–882.

Ogryzko, N. V., Renshaw, S. A. and Wilson, H. L. (2014a). The IL-1 family in fish: Swimming through the muddy waters of inflammasome evolution. Dev. Comp. Immunol. 46, 53–62.

Ogryzko, N. V., Hoggett, E. E., Solaymani-Kohal, S., Tazzyman, S., Chico, T. J. A., Renshaw, S. A. and Wilson, H. L. (2014b). Zebrafish tissue injury causes upregulation of interleukin-1 and caspase-dependent amplification of the inflammatory response. Dis. Model. Mech. 7, 259–264.

Rébé, C. and Ghiringhelli, F. (2020). Interleukin-1β and Cancer. Cancers (Basel). 12,.

Reis, M. I. R., Vale, A., Pereira, P. J. B., Azevedo, J. E. and Santos, N. M. S. (2012). Caspase-1 and IL-1 b Processing in a Teleost Fish. 7,.

Rider, P., Voronov, E., Dinarello, C. A., Apte, R. N. and Cohen, I. (2017). Alarmins: Feel the Stress. J. Immunol. 198, 1395–1402.

Rivers-Auty, J., Daniels, M. J. D., Colliver, I., Robertson, D. L. and Brough, D. (2018). Redefining the ancestral origins of the interleukin-1 superfamily. Nat. Commun. 9, 1–12.

Suzuki, T., Hashimoto, S. I., Toyoda, N., Nagai, S., Yamazaki, N., Dong, H. Y., Sakai, J., Yamashita, T., Nukiwa, T. and Matsushima, K. (2000). Comprehensive gene expression profile of LPS-stimulated human monocytes by SAGE. Blood 96, 2584–2591.

Szymczak, A. L., Workman, C. J., Wang, Y., Vignali, K. M., Dilioglou, S., Vanin, E. F. and Vignali, D. A. A. (2004). Correction of multi-gene deficiency in vivo using a single “self-cleaving” 2A peptide-based retroviral vector. Nat. Biotechnol. 22, 589–594.

Takacs, L., Kovacs, E. J., Smith, M. R., Young, H. A. and Durum, S. K. (1988). Detection of IL-1 alpha and IL-1 beta gene expression by in situ hybridization. Tissue localization of IL-1 mRNA in the normal L Takács, E J Kovacs, M R Smith, H A Young and S K Why The JI ? Submit online. • Rapid Reviews ! 30 days * from submission. J. Immunol. 141, 3081–2095.

Tapia, V. S., Daniels, M. J. D. Palazón-Riquelme, P., Dewhurst, M., Luheshi, N. M., Rivers-Auty, J., Green, J., Redondo-Castro, E., Kaldis, P., Lopez-Castejon, G., et al. (2019). The three cytokines IL-1β, IL-18, and IL-1α share related but distinct secretory routes. J. Biol. Chem. 294, 8325–8335.

Thornberry, N. A., Bull, H. G., Calaycay, J. R., Chapman, K. T., Howard, A. D., Kostura, M. J., Miller, D. K., Molineaux, S. M., Weidner, J. R., Aunins, J., et al. (1992). A novel heterodimeric cysteine protease is required for interleukin-1β processing in monocytes. Nature 355, 242–244.

Tsuchiya, K., Nakajima, S., Hosojima, S., Thi Nguyen, D., Hattori, T., Manh Le, T., Hori, O., Mahib, M. R., Yamaguchi, Y., Miura, M., et al. (2019). Caspase-1 initiates apoptosis in the absence of gasdermin D. Nat. Commun. 10,.

Vojtech, L. N., Scharping, N., Woodson, J. C. and Hansen, J. D. (2012). Roles of inflammatory caspases during processing of zebrafish interleukin-1β in Francisella noatunensis infection. Infect. Immun. 80, 2878–2885.

Young, P. R. and Sylvester, D. (1989). Cloning of rabbit interleukin-1β: differential evolution of IL-1α and IL-1β proteins. Protein Eng. Des. Sel. 2, 545–551.

