## Supplementary figures 1&2 and Tables for "Revisiting the origin of interleukin 1 (IL-1) based on biological activities of IL-1 in anamniotes and their sub-functionalization in amniotes" (from another version of this preprint: version 1, 2022-02-15)

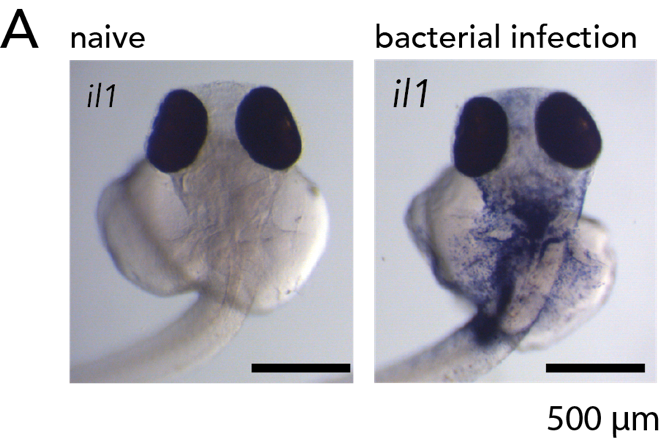

Supplementary figure 1:

il-1 expression detected by in situ hybridization with il-1 antisense probe in uninfected medaka larvae (left) and in larvae injected with *E. coli* as 6 day old embryos

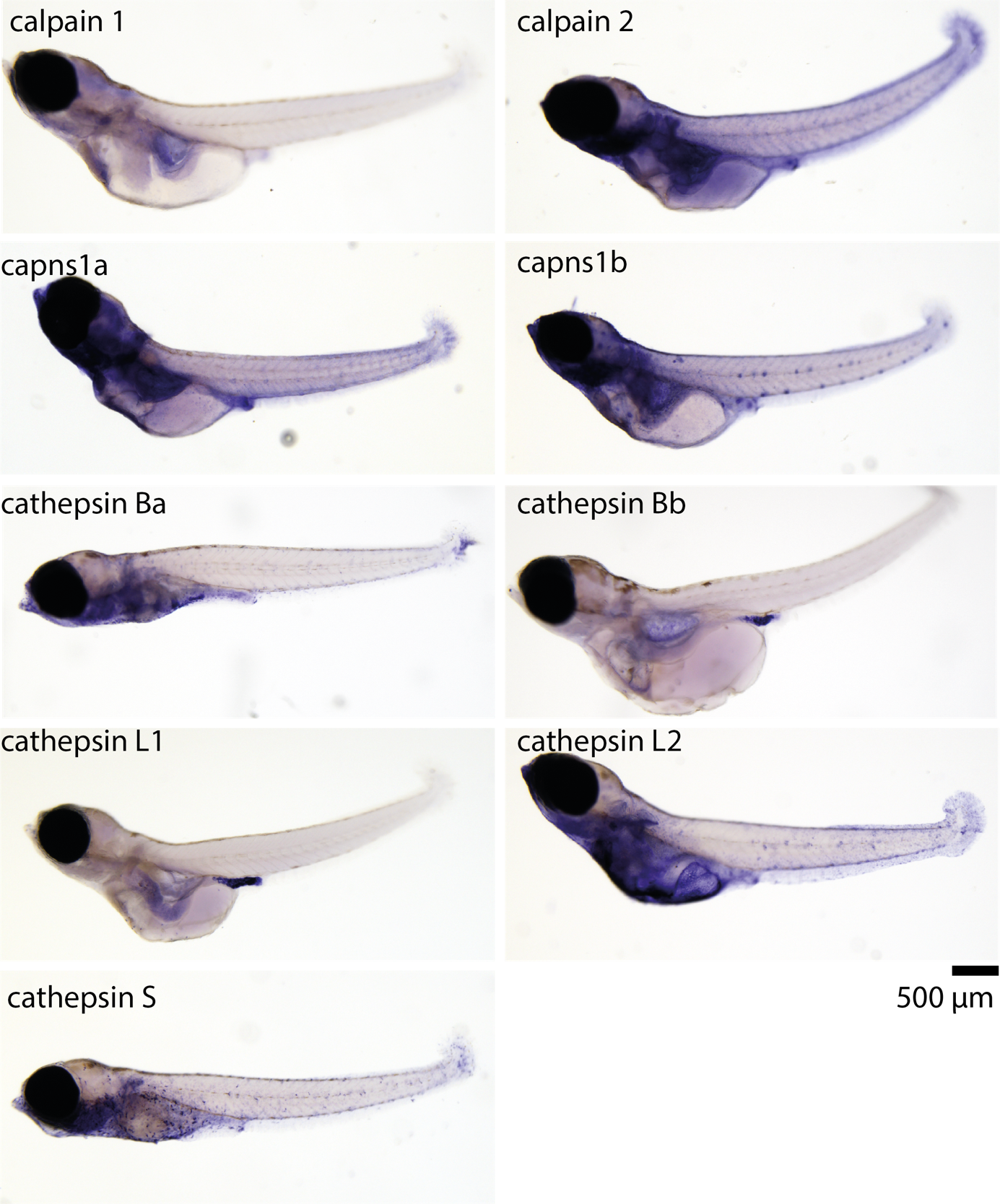

Supplementary figure 2: In situ hybridization showing expression of the indicated calpain and cathepsin genes in 10 dpf medaka larvae.

**Supplementary Table 1: Gene-specific in situ hybridization probes used in this study**

| **Gene** | **Ensembl ID/Accession Number** | **Nucleotides of transcript** |
| --- | --- | --- |
| il-1 | ENSORLG00000000217 | 19-777 |
| cathepsin Ba | ENSORLG00000011080 | 629-1069 |
| cathepsin Bb | ENSORLG00000017347 | 241-966 |
| cathepsin S | ENSORLG00000015733 | 107-1096 |
| cathepsin L1 | ENSORLG00000012518 | 913-1670 |
| cathepsin L | ENSORLG00000020801 | 124-937 |
| cathepsin S | ENSORLG00000015733 | 59-965 |
| calpain1 | ENSORLG00000011738 | 1002-1838 |
| calpain2 | ENSORLG00000020892 | 1120-1905 |
| capns1a | ENSORLG00000000369 | 554-1162 |
| capns1b | ENSORLG00000000490 | 56-649 |

**Supplementary Table 2: List of IL-1 protein sequences used for sequence alignment (alphabetically ordered)**

| **Species** | **Species name in Alignment** | **Gene** | **Ensembl/NCBI identifier** | **Size** |
| --- | --- | --- | --- | --- |
| Alligator mississippiensis | **alligator** | **IL-1ß** | [KYO25690.1](https://www.ncbi.nlm.nih.gov/protein/KYO25690.1?report=genbank&log$=protalign&blast_rank=6&RID=PDEDY2VP01T) | 269 aa |
| Ursus americanus | **bear** | **IL-1α** | ENSUAMT00000036710.1 | 270 aa |
| Ursus americanus | **bear** | **IL-1ß** | ENSUAMT00000036707.1 | 269 aa |
| Gallus gallus | **chicken** | **IL-1ß** | NP_989855 | 270 aa |
| [Chrysemys picta bellii](https://www.ncbi.nlm.nih.gov/taxonomy/8478) | **turtle** | **IL-1ß** | [XP_005279317.1](https://www.ncbi.nlm.nih.gov/protein/XP_005279317.1?report=genbank&log$=protalign&blast_rank=1&RID=PG76Y3V701N) | 267 aa |
| Cyprinus carpio | **carp** | **IL-1ßlike** | ENSCCRT00000074741.1 | 266 aa |
| Cyprinus carpio | **carp** | **IL-1ßlike** | ENSCCRT00000015599.1 | 270 aa |
| Anas platyrhynchos platyrhynchos | **duck** | **IL-1ß** | XP_038022797 | 269 aa |
| Epinephelus lanceolatus | **grouper** | **IL-1ß** | [XP_033475983.1](https://www.ncbi.nlm.nih.gov/protein/XP_033475983.1?report=genbank&log$=protalign&blast_rank=2&RID=R037YR6401N) | 320 aa |
| Homo sapiens | **human** | **IL-1α** | ENST00000263339.4 | 269 aa |
| Homo sapiens | **human** | **IL-1ß** | ENST00000263341.7 | 269 aa |
| Myotis lucifugus | **bat** | **IL-1α** | ENSMLUT00000000165.2 | 244 aa |
| Myotis lucifugus | **bat** | **IL-1ß** | ENSMLUT00000009938.2 | 269 aa |
| Lynx canadensis | **lynx** | **IL-1α** | ENSLCNT00005025995.1 | 272 aa |
| Lynx canadensis | **lynx** | **IL-1ß** | ENSLCNT00005025973.1 | 263 aa |
| Siniperca chuatsi | **mandarin fish** | **IL-1ß** | AAV65041 | 275 aa |
| Oryzias latipes | **medaka** | **IL-1ß** | ENSORLG00000000217 | 251 aa |
| Mus musculus | **mouse** | **IL-1α** | ENSMUST00000028882.2 | 337 aa |
| Mus musculus | **mouse** | **IL-1ß** | ENSMUSG00000027398 | 448 aa |
| Didelphis | **opossum** | **IL-1ß** | ENSMODT00000017586.3 | 286 aa |
| Didelphis | **opossum** | **IL-1α** | ENSMODT00000017594.3 | 283 aa |
| Sus Scofa | **pig** | **IL-1α** | ENSSSCT00000008863.3 | 278 aa |
| Sus Scofa | **pig** | **IL-1ß** | [ENSSSCT00000008860.4](http://www.ensembl.org/Sus_scrofa/Transcript/Summary?db=core;g=ENSSSCG00000008087;r=AEMK02000612.1:87520-104056;t=ENSSSCT00000008860) | 272 aa |
| Columba livia | **pigeon** | **IL-1ß** | [NP_001269753.1](https://www.ncbi.nlm.nih.gov/protein/NP_001269753.1?report=genbank&log$=protalign&blast_rank=1&RID=R2R6NP5Z013) | 294 aa |
| Ornithorhynchus anatinus | **platypus** | **IL-1ß** | ENSOANT00000016715.3 | 269 aa |
| Coturnix coturnix | **quail** | **IL-1ß** | ACY72337 | 262 aa |
| Salmo salar | **salmon** | **il--1ß** | [XP_014025954.1](https://www.ncbi.nlm.nih.gov/protein/XP_014025954.1?report=genbank&log$=protalign&blast_rank=1&RID=R2TZFDYT013) | 251 aa |
| Scyliorhinus canicula | **shark** | **IL-1ß** | CAC09435.1 | 272 aa |
| Physeter catodon | **sperm whale** | **IL-1α** | ENSPCTT00005012000.1 | 258 aa |
| Physeter catodon | **sperm whale** | **IL-1ß** | [ENSPCTT00005011958.1](http://www.ensembl.org/Physeter_catodon/Transcript/Summary?db=core;g=ENSPCTG00005007680;r=12:12768913-12774658;t=ENSPCTT00005011958) | 301 aa |
| Gasterosteus aculeatus | **stickleback** | **IL-1ß** | [CCV66729.1](https://www.ncbi.nlm.nih.gov/protein/CCV66729.1?report=genbank&log$=protalign&blast_rank=1&RID=R2S5DM25016) | 369 aa |
| Xenopus laevis | **xenopus laevis** | **IL-1ß-like** | [CAC85480.1](https://www.ncbi.nlm.nih.gov/protein/CAC85480.1?report=genbank&log$=protalign&blast_rank=7&RID=PDERA94401T) | 254 aa |
| Xenopus tropicalis | **xenopus tropicalis** | **IL-1ß** | [XP_002932618.1](https://www.ncbi.nlm.nih.gov/protein/XP_002932618.1?report=genbank&log$=protalign&blast_rank=4&RID=PDEMJW7M01T) | 267 aa |
| Danio rerio | **zebrafish** | **IL-1ß** | ENSDARG00000098700 | 280 aa |
